# The Neuroprotective Beta Amyloid Hexapeptide Core Reverses Deficits in Synaptic Plasticity in the 5×FAD APP/PS1 Mouse Model

**DOI:** 10.1101/2020.03.17.995191

**Authors:** Kelly H. Forest, Ruth Taketa, Komal Arora, Cedomir Todorovic, Robert A. Nichols

**Affiliations:** Department of Cell & Molecular Biology, John A. Burns School of Medicine, University of Hawai’i at Manoa, Honolulu, HI, USA

## Abstract

Alzheimer’s disease (AD) is the most common cause of dementia in the aging population. Evidence implicates elevated soluble oligomeric Aβ as one of the primary triggers during the prodromic phase leading to AD, effected largely via hyperphosphorylation of the microtubule-associated protein tau. At low, physiological levels (pM-nM), however, oligomeric Aβ has been found to regulate synaptic plasticity as a neuromodulator. Through mutational analysis, we found a core hexapeptide sequence within the N-terminal domain of Aβ (N-Aβcore) accounting for its physiological activity, and subsequently found that the N-Aβcore peptide is neuroprotective. Here, we characterized the neuroprotective potential of the N-Aβcore against dysfunction of synaptic plasticity assessed in *ex vivo* hippocampal slices from 5×FAD APP/PS1 mice, specifically hippocampal long-term potentiation (LTP) and long-term depression (LTD). The N-Aβcore was shown to reverse impairment in synaptic plasticity in hippocampal slices from 5×FAD APP/PS1 model mice, both for LTP and LTD. The reversal by the N-Aβcore correlated with alleviation of downregulation of hippocampal AMPA-type glutamate receptors in preparations from 5×FAD mice. The action of the N-Aβcore depended upon a critical di-histidine sequence and involved the PI3 kinase pathway via mTOR. Together, the present findings indicate that the non-toxic N-Aβcore hexapeptide is not only neuroprotective at the cellular level but is able to reverse synaptic dysfunction in AD-like models, specifically alterations in synaptic plasticity.

## Introduction

Alzheimer’s disease (AD) is clinically characterized by impairments in cognitive memory and function. Loss of critical pre- and post-synaptic markers have been reported for postmortem AD brain tissue [1,2], suggesting that AD-related cognitive impairments are based, in large part, on synaptic dysfunction and loss. Additionally, accumulating evidence shows a strong link between excess soluble oligomeric amyloid-β (Aβ) and synaptic dysfunction in AD [3-5]. Cognitive decline and synaptic plasticity deficits are reported to occur prior to the accumulation of Aβ plaques and tau neurofibrillary tangles in prodromic phases leading to AD [6], supporting the idea that synaptic dysfunction and mild cognitive impairment are early events driven by soluble oligomeric Aβ rising to abnormally high levels years prior to AD diagnosis.

Synaptic dysfunction and eventual degeneration lead to abnormal synaptic transmission and impaired long-term potentiation (LTP) and/or long-term depression (LTD), which are important in synaptic plasticity and learning and memory. Pathological levels (high nM to μM) of Aβ have been shown to inhibit LTP-induction [3,7,8] and enhance LTD [9,10] in the hippocampus. On the other hand, low physiological levels (pM) of Aβ was found to enhance LTP and memory, indicating a hormetic effect of Aβ on synaptic plasticity [11-13].

Dysregulation of synaptic plasticity in AD pathogenesis involves altered regulation of NMDA-type and AMPA-type glutamate receptors. In addition to mediating Aβ-induced excitotoxicity, NMDA receptors can be depressed by Aβ at high concentrations [14], inducing LTD [15,16] as a consequence of subsequent downstream AMPA receptor internalization [15,16] and dendritic spine loss [16].

We have shown that at low concentration (pM-nM) the N-terminal Aβ fragment comprising amino acids 1-15/16 of the Aβ sequence, an endogenous peptide cleaved from Aβ via α-secretase [17], is more effective as a neuromodulator than full-length Aβ1-42, stimulating receptor-linked increases in neuronal Ca^2+^, enhancing synaptic plasticity and enhancing contextual fear memory [13]. The Aβ1-16 peptide sequence corresponds to the C-terminal 16 amino acid sequence in soluble amyloid precursor protein-α (sAPP-α), referred to as the CTα16, which has also been shown to enhance synaptic plasticity [18]. An essential core sequence comprising amino acids 10-15 of Aβ (N-Aβcore) was identified as the active region of the N-terminal Aβ fragment and was further shown to protect against Aβ-induced neuronal toxicity [19]. Here, we aimed to better understand the neuroprotective mechanism of the N-Aβcore on synaptic plasticity. We investigated whether the N-Aβcore could rescue LTP and LTD dysfunction resulting from prolonged, elevated levels of Aβ in an APP/PS1 transgenic mouse model harboring mutations found in familial Alzheimer’s disease (FAD), while assessing the impact on AMPA-type glutamate receptor expression in reference to the neuroprotective action of the N-Aβcore in Aβ-synaptotoxicity.

## Materials and methods

### Transgenic mice and cannulation surgery

All animal handling, surgery, use and euthanasia were performed under an approved IACUC protocol (11-1219-6 / 16-2282-2), compliant with NIH and Society for Neuroscience guidelines for use of vertebrate animals in neuroscience research. The human APP/PSEN1 mouse line, 5×FAD (Tg6799), on the B6.SJL background (B6SJL-Tg(APPSwFlLOn,PSEN1*M146L*L286V) 6799Vas/Mmjax; obtained from JAX stock #006554, MMRRC034840 hemizygous) was used as a well characterized model for Aβ-based pathology and neurodegeneration [20], along with age-matched control (B6.SJL background) mice (MMRRC034840 Non-carrier). Mice at 7- to 8-months of age of both sexes (weight range: 28-35g), obtained from established in-house colonies and housed in ventilated enrichment cages in the John A. Burns School of Medicine AAALAC-accredited Vivarium with *ad libitum* access to food and water, were used at roughly equal numbers, this age range selected for displaying pronounced LTP deficits in the transgenic line. Inclusion/exclusion criteria were based on animal health.

For bilateral cannulation and injection, the following protocol was employed (as per ref 19). Cannulation into the dorsal aspects of both hippocampi of the 5×FAD mice at 7- to 8-months of age of both sexes was performed under full anesthesia (general: 1.2% Avertin; local at site: lidocaine) using stereotaxis (coordinates: −1.5mm anteroposterior; ±1mm lateral; −2mm depth). After the brief surgical procedure and recovery (full righting reflex), mice were subsequently housed in sound-isolated, ventilated hotels prior to peptide injection one week later. On the day of microinjection (morning), sterile saline or 500 nM N-Aβcore peptide was administered bilaterally through the cannulae in the 5×FAD mice via microinjectors over 30s (0.5μL/side) and the mice were returned to their cages in the mouse hotel. Hippocampi were collected from euthanized mice 24 h after the bilateral microinjection of the saline or peptide, and lysates extracted from the hippocampi were prepared for immunoblot analysis (30 µg each). Euthanasia was performed under an approved IACUC protocol (11-1219-6 / 16-2282-2). This study was not preregistered and followed ARRIVE guidelines.

### Preparation of Aβ peptides

Solutions of Aβ_1-42_ (American Peptide; Anaspec) were prepared from aqueous stock solutions, followed by bath sonication. This preparation of full-length Aβ was previously shown to exist predominantly in the oligomeric state [see 13]. The N-Aβcore peptide (synthesized and purified to Peptide 2.0) was prepared from aqueous stock solutions.

### Extracellular field potential recordings in hippocampal slices

Hippocampal slices were prepared from 7- to 8-month-old 5×FAD (Tg6799) or B6.SJL (control mice) (as per ref 13). Cervical dislocation and decapitation were performed under an approved IACUC protocol (11-1219-6 / 16-2282-2), compliant with NIH and Society for Neuroscience guidelines for use of vertebrate animals in neuroscience research. Brains were removed into ice-cold artificial cerebral spinal fluid (aCSF) consisting of 130mM NaCl, 3.5 mM KCl, 10mM glucose, 1.25mM NaH_2_PO_4_, 2.0mM CaCl_2_, 1.5mM MgSO_4_, and 24mM NaHCO_3,_ bubbled in 95% O_2_/5% CO_2_. Transverse brain slices of 400μm were obtained using a Leica vibrating microtome (Leica, VT1200S) and quickly transferred to fresh ice-cold aCSF for hippocampi isolation. Extracted hippocampi slices were incubated in bubbled aCSF in a holding chamber for 30 mins at room temperature (23°C) after which the holding chamber was transferred to a 32°C water bath for 1 h. The chamber was then removed from the water bath and placed at room temperature for another 1 h prior to recording. The slices were subsequently transferred to a recording chamber and perfused at 3mL/min with aCSF (bubbled with 95% O_2_/5% CO_2_) at 32°C. The Schaffer collateral fibers were stimulated at a frequency 0.1 Hz using a bipolar stimulating electrode and CA1 field excitatory postsynaptic potentials (fEPSPs) were recorded with a glass electrode filled with 3M NaCl (resistance 1-1.5 MΩ). Basal synaptic transmission was assessed by comparing stimulus strength against fEPSP slope to generate input/output (I/O) curves. A minimum of 20 min baseline stimulation was then performed, recording every minute. The baseline and stimulus current were adjusted during this period so that fEPSP stabilized at 30-40% of maximum.

LTP was induced by a 3-theta-burst stimulation (TBS) protocol, where each burst consisted of 4 pulses at 100 Hz with a 200-ms interburst interval. LTD was induced using a low frequency stimulation (LFS) protocol, consisting of a 1Hz single pulse stimulus (900 pulses for 15 min). TBS and LFS were administered after a 20-min baseline recording period for aCSF alone or a 35-min baseline recording period in aCSF (15 min) followed by inclusion of N-Aβcore in the absence or presence of tested reagents (20 min). For the latter, TBS and LFS were administered in the presence of N-Aβcore in the absence or presence of tested reagents.

### Immunoblot analysis

#### Hippocampi injected with N-Aβcore

Hippocampi removed from euthanized 8-month-old 5×FAD mice previously bilaterally injected with saline or N-Aβcore were homogenized with 200-250 μL of Pierce I.P Lysis Buffer (ThermoFisher Scientific, # 87788, lot# MJ162614) with 1x Halt Protease and Phosphatase Inhibitor Cocktail (ThermoFisher Scientific, # 78441, lot# SF248390). The homogenates were centrifuged at 18,000*g* for 20 min at 4°C and the supernatant was collected. The total amount of protein was quantified by a Pierce™ BCA Protein Assay Kit (ThermoFisher Scientific, # 23225).

For each condition, gel sample buffer (4x; ThermoFisher Scientific, # B0007, lot # 1920132) and reducing agent (10x; ThermoFisher Scientific, # B0009, lot # 1901009) were added to diluted SDS-solubilized protein samples for a final protein concentration of 2 μg/μL. The samples were boiled at 95°C for 10 min, immediately cooled on ice and then centrifuged. Equal amounts of protein were subjected to electrophoretic separation on a 4-20% Tris-Glycine polyacrylamide gel (ThermoFisher Scientific, #XP04200), transferred to Nitrocellulose membrane (LI-COR, # 92631092) via the iBlot2 semidry system (ThermoFisher). Recovered blots were incubated in primary antibody (see below) overnight at 4°C. The transfer blots were washed 3x (10 min each wash) in 0.1% Tween-20 in Tris-buffered saline (TBS) and incubated in the appropriate IR-dye-conjugated secondary antibody (LI-COR Biosciences) for 1h. An Odyssey IR imaging system (LI-COR Biosciences, Lincoln, NE) was used for signal detection. Analysis was performed via Image Studio v5.2.5 software (LI-COR Biosciences).

The following primary antibodies were used for detection and normalization, respectively: anti-pAMPAR1 rabbit monoclonal antibody (pS831-GluA1; Cell Signaling Technology; RRID:AB_2799873) and anti-AMPAR1 rabbit monoclonal antibody (GluA1; Cell Signaling Technology; RRID:AB_2732897), both at 1:1000 and anti-beta actin mouse monoclonal antibody (Sigma-Aldrich; RRID:AB_476697) at 1:10,000 dilution.

### Cultured hippocampal neurons

Hippocampal neuron cultures were prepared as described [21] from neonatal mouse pups (0-2d old; one litter (6-10 mice)/preparation) from female breeders housed in the John A. Burns School of Medicine AAALAC-accredited Vivarium of either gender (equivalent numbers) obtained from established colonies of wild-type B6.SJL (background) mice. All animal procedures (handling; euthanasia) followed an IACUC-approved protocol (16-2282-2), compliant with NIH and Society for Neuroscience guidelines for use of vertebrate animals in neuroscience research. Following rapid decapitation, brains were removed from the mice into ice-cold Neurobasal A medium (NB) containing B-27 supplement, 5% fetal bovine serum (FBS) and Gentamicin. Hippocampi were then dissected under a stereomicroscope (Olympus SZ61). The isolated hippocampi were treated with papain (Worthington, LS003126, Lot # 35N16202) in Hanks buffer with 10mM cysteine at 37°C for 15mins. The partially digested tissue was washed by centrifugation in NB plus FBS. The cells were dissociated using sequential trituration (polished Pasteur pipettes of decreasing diameter) and collected by low-speed centrifugation. The dissociated cells were pre-plated in standard tissue culture dishes to remove adherent non-neuronal cells (glia; fibroblasts) for 10-15 mins. The neuron-enriched preparation was diluted to 1×10^5^ cells/mL and then plated into poly-D-lysine-coated 24-well dishes in NB plus serum and Gentamicin. The cultures were washed with Neurobasal A medium containing B27 and Gentamicin to remove the serum and then cultured in this media for 7-10 days prior to treatment with Aβ, N-Aβcore or the combination for an additional 7 days.

### qPCR

RNA was extracted from treated cultured hippocampal neurons using the PureLink® RNA Mini Kit (Ambion, Life Technologies, #12183025) as per the manufacturer’s protocol. Genomic DNA contamination was eliminated the RNA preparation by digesting with RNase-free DNase (Qiagen, #79254). The iScript™ cDNA Synthesis Kit (Bio-Rad) was used to synthesize cDNA. The expression levels of various genes were then determined using SYBR green via qPCR (Bio-Rad iCycler iQ™ Multicolor Real-Time PCR Detection System) using the primers listed in the accompanying table (Table 1). Cycling conditions were as follows: 95 °C for 15 min, followed by 40 cycles of 94 °C for 15 s and 60 °C for 60 s and the fold-changes in the variously treated samples compared to untreated (vehicle control) samples were calculated after normalizing to GAPDH gene expression.

**Table 1.**
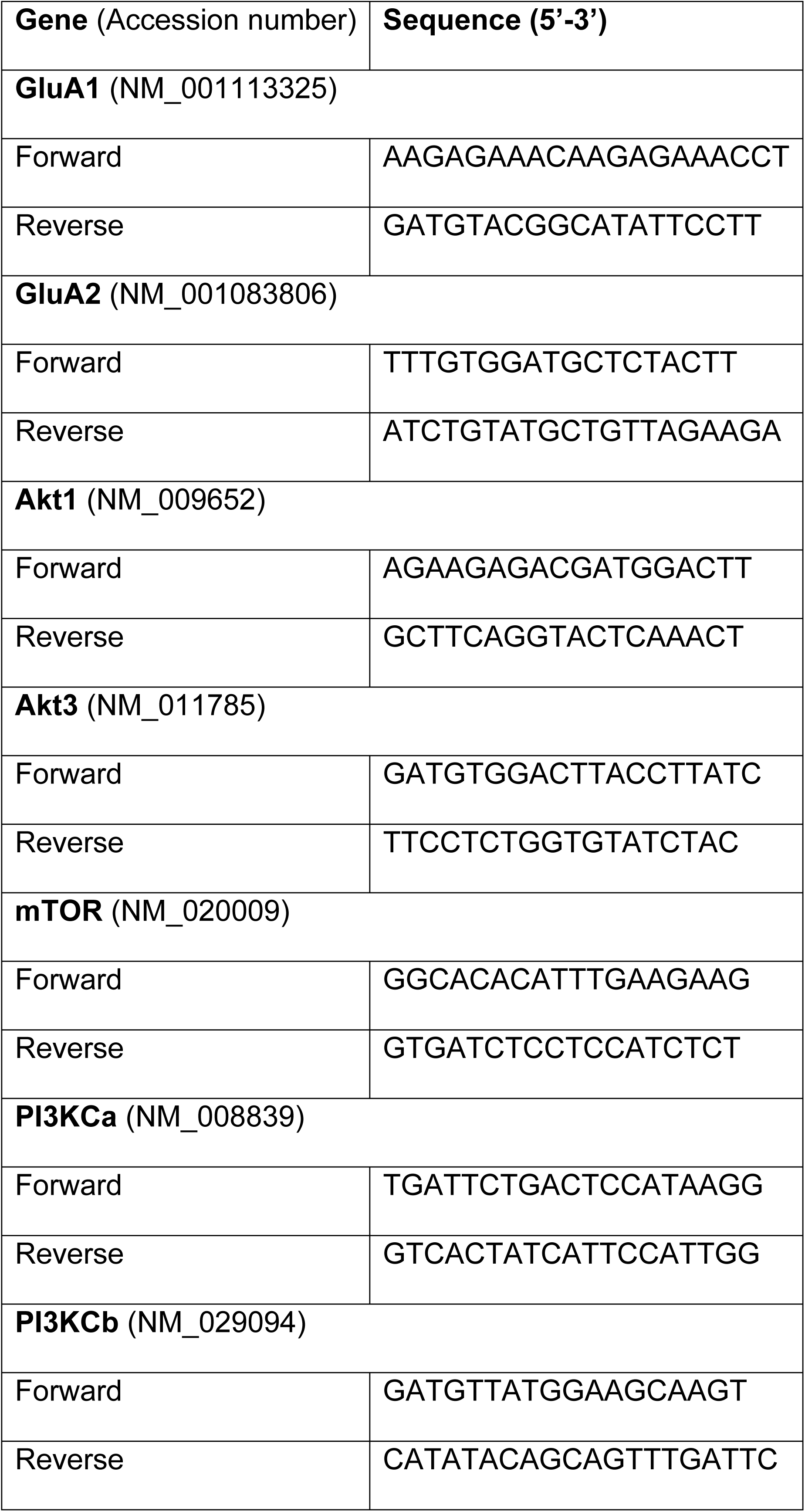

### Data and statistical analysis

Treatment and units were randomized as to order for all assays and experiments. Biological replicates were based on independent samples (*n*). All experiments were repeated at least three times unless otherwise noted. After testing for normality, multiple comparisons of the data were made using one-way ANOVA with Bonferroni or Tukey’s post hoc tests, as indicated. Paired comparison was made using Student’s *t*-tests. *P*-values <0.05 were considered the minimum for significance (as rejection of the null hypothesis). Unless otherwise noted, data were analyzed and graphed with GraphPad Prism 5 (GraphPad v5.0b; RRID:SCR_002798) using the appropriate statistical tests.

## Results

### The N-Aβcore reversed LTP deficits induced by pathological levels of full-length Aβ

We have previously shown that the N-terminal Aβ fragment (Aβ_1-15/16_) enhances synaptic plasticity and contextual fear memory while protecting against Aβ-linked synaptic impairment [13]. Considering the evidence that the N-Aβcore accounts for the neuromodulatory activity of the N-terminal Aβ fragment, we assessed whether the N-Aβcore is capable of reversing Aβ-linked synaptic dysfunction in an *ex vivo* model. We utilized hippocampal slices from a mutant APP/PS1 transgenic AD-like endophenotype mouse model (5×FAD) and their wild-type counterparts (B6.SJL) to examine synaptic transmission. Basal synaptic transmission at the Schaffer collateral-CA1 synapses represented by input-output curves shows that the fEPSP slopes versus stimulus strength for both the 5×FAD and B6.SJL mice were comparable (Fig 1A), ruling out any issues with regard to impact of the Aβ fragment peptides on baseline synaptic strength. Interestingly, treatment with the N-Aβcore during baseline recordings induced increases in baseline synaptic transmissions for both 5×FAD and B6.SJL but was only significant in the B6.SJL slices (Fig 1B; average increase relative to untreated controls: 110% ± 2% s.d.).

**Fig. 1.**
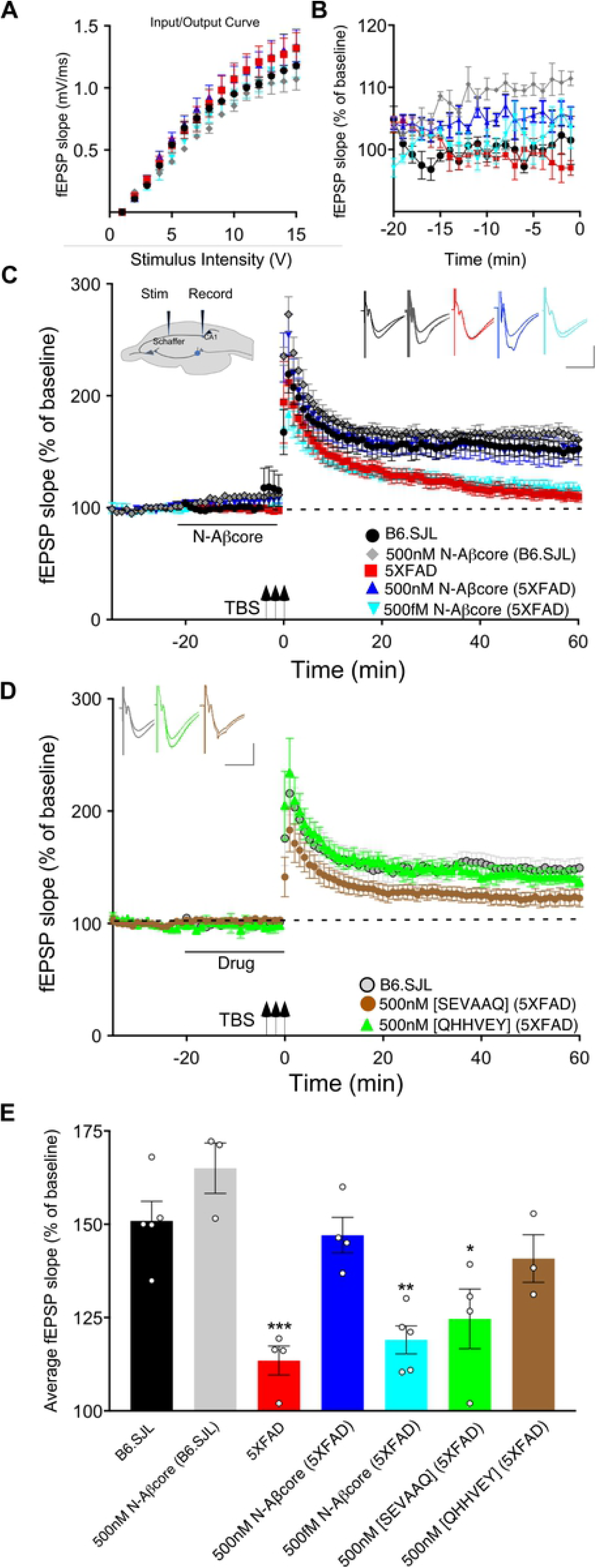
The N-Aβcore reverses Aβ-induced synaptic impairment in APP/PSEN1 mutant mouse hippocampal slices. Input/Output curves (**A)** plotting average fEPSP slopes versus stimulus strength before treatment and baseline recordings (**B**) during drug treatment versus aCSF perfusion alone prior to stimulation for: control aCSF in B6.SJL slices (black, *n*=5), 5×FAD slices (red, *n*=4), 500nM N-Aβcore in B6.SJL slices (grey, *n*=3) and 5×FAD slices (blue, *n*=4), plus 500fM N-Aβcore in 5×FAD slices (cyan, *n*=5). No statistically significant differences in the input/output curves (**A**) were found (p>0.05, one-way repeated measures ANOVA). Averaged (last 10mins**) baseline data (**B**) were analyzed by one-way ANOVA (F_(4,16)_= 7.53, p=0.0013). Bonferroni post hoc tests revealed significant enhancement of baseline fEPSPs by 500nM N-Aβcore on B6.SJL slices compared to aCSF-perfused B6.SJL slices (p=0.006), aCSF perfused 5×FAD slices (p= 0.002), and by 500fM N-Aβcore on 5×FAD slices (p= 0.038). **C** TBS-induced LTP for B6.SJL slices perfused with control aCSF (black, *N*=5) or 500nM N-Aβcore (grey, *N*=3) and 5×FAD slices with aCSF (red, *N*=4), 500nM N-Aβcore (blue, *N*=4) and 500fM N-Aβcore (cyan, *N*=5). Inset in **c**, diagram of hippocampal slice stimulation of the Schaffer collaterals (Stim) and recording fEPSPs in CA1 (Record). **D** TBS-induced LTP for 5×FAD slices treated with 500nM [SEVAAQ] (green, *n*=5) or 500nM [QHHVEY] (brown, *n*=3) substituted N-Aβcore peptides, and, as a reference, B6.SJL perfused with control aCSF (gray circles, *n*=5, data from **C**). **E** Quantification of average fEPSP slope values for 50-60 min posTBS. All groups shown in panels in **C** and **D** were analyzed by one-way ANOVA (F_(6,21)_= 11.33, p< 0.0001). Bonferroni post hoc tests: differences found for 5×FAD slices (p= 0.0012), 500fM N-Aβcore in 5×FAD slices (p= 0.0036), and 500nM [SEVAAQ] in 5×FAD slices (p=0.041) when compared to aCSF-perfused (control) B6.SJL slices. Application of 500nM N-Aβcore in 5XFAD slices reversed the LTP deficit in the 5XFAD slices (p<0.0005). Horizontal bars in **C** and **D** indicate the period (20min) of drug delivery for the color-coded conditions as indicated. Arrows indicate timing of TBS in **C** and **D**. Color-coded insets showing examples of fEPSPs (baseline vs. LTP). All data are means ± s.e.m. *N* values represent independent experiments (1 slice/mouse). Inset calibration: horizontal, 10ms; vertical, 0.4 mV. *p< 0.05; **p<0.005, ***p<0.0005.

To assess sustained changes in synaptic plasticity, we used a 3x-TBS stimulation protocol at the Schaffer collaterals to measure LTP in the CA1 region (see cross-sectional diagram of the hippocampus in Fig 1A, inset). LTP showed a trend toward enhancement for the N-Aβcore-treated B6.SJL slices as compared to untreated slice, though it was not significant (Figs 1C&1E). Consistent with previous findings [22], LTP in the 5×FAD slices was substantially reduced compared to that observed for slices from B6.SJL (Figs 1C&1E; 24.2% of control), dropping to near baseline at 60mins post-TBS. By contrast, prestimulation treatment with 500nM N-Aβcore restored LTP in the 5×FAD slices to the level seen for the wild-type slices (Figs 1C&1E; 107% ± 44% s.d. of control). These findings demonstrate that the N-Aβcore can reverse LTP deficits induced from prolonged exposure to pathological levels of Aβ, while modestly enhancing basal synaptic transmission.

### The N-Aβcore reversed full-length Aβ-linked downregulation of hippocampal AMPA-type glutamate receptors

Regulation of synaptic expression of AMPA-type glutamate receptors (AMPARs) has been shown to underlie LTP [23,24]. As downregulation of AMPARs is linked to the impairment in hippocampal LTP in APP/PS1 mice [15], the impact *in vitro* and *in vivo* of the N-Aβcore on the regulation of hippocampal AMPARs was assessed.

Utilizing hippocampal neuronal cultures derived from the background wild-type mice, B6.SJL, and treated with exogenous full-length Aβ (Aβ_42_) for 7 days, the N-Aβcore was shown to alleviate the downregulation of AMPAR1 (GluA1) transcript expression assessed via qPCR in this *in vitro* Aβ toxicity model (Fig 2A; 22% ± 0.8% s.d. of Aβ_42_ condition). There was no significant impact of the N-Aβcore on the modest downregulation of AMPAR2 (GluA2) transcript expression.

**Fig. 2.**
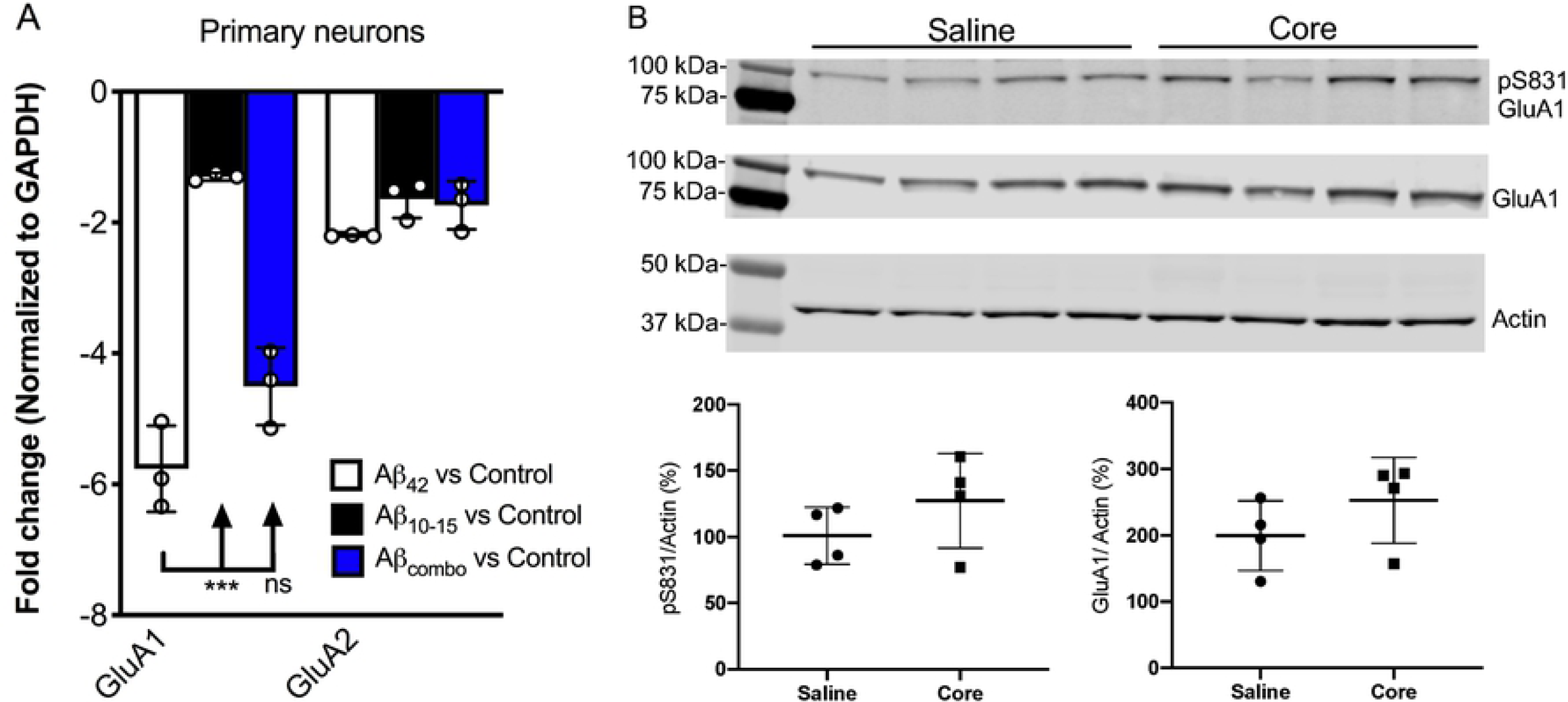
N-AβCore normalizes AMPAR1 expression in an *in vitro* Aβ neurotoxicity model of primary hippocampal neurons and *in vivo* in hippocampi of 5×FAD APP/PS1 mice. **A** Normalized average expression of AMPA receptor GluA1 and GluA2 transcripts in B6.SJL mouse hippocampal neuron cultures exposed to full-length Aβ_1-42_ (Aβ_42_), the N-Aβcore (Aβ_10-15_) or the combination (combo), as measured via qPCR. Data are means ± s.d. *N*=3 cultures for each condition. ***p=0.000001 (Bonferroni post hoc tests for one-way ANOVA comparison between Aβ_42_ and Aβ_10-15_). There was no statistical difference (ns) between expression of GluA1 for Aβ_42_ and Aβ_42_ plus Aβ_10-15_ (combo), or for any condition for GluA2. **B** Expression of pAMPAR1 (pGluA1) or total AMPAR1 (GluA1) in the hippocampi of 5xFAD mice injected with sterile saline or N-Aβcore (Core), as measured using western immunoblot. *N*=4 mice for each condition. Total GluA1: p=0.247; pGluA1: p=0.252. (Student’s *t*-tests for comparison of saline vs. N-Aβcore).

Proteins solubilized from hippocampi isolated from 5×FAD mice bilaterally injected with the N-Aβcore or saline vehicle were assayed for changes in *in vivo* expression of hippocampal AMPAR1 (GluA1) via immunoblot analysis. Total GluA1 levels in the hippocampi were increased with exposure to the N-Aβcore as compared to saline-injected controls (Fig 2B; 248% ± 67% s.d.). The increase in relative pAMPAR1 (pS831) in the hippocampi exposed to N-Aβcore was accounted by the increase in total GluA1 (Fig 2B; 127% ± 35% s.d.). Together, these findings indicate that reversal of the LTP impairment in the 5×FAD APP/PS1 by the N-Aβcore involves regulation of AMPAR expression.

### Structure-function and concentration-dependence of the N-Aβcore in reversing LTP impairment in hippocampal slices from 5×FAD APP/PS1 mice

We also tested for basic concentration-dependence of the N-Aβcore in reversing LTP impairment in the 5×FAD mouse hippocampal slices. Low concentration (<pM) of the N-Aβcore showed no difference on LTP compared to control 5×FAD slices (Fig 1C & 1E; 107% ± 7% s.d. compared to 5×FAD).

Through Aβ-interacting receptor-linked Ca^2+^ and neurotoxicity assays, we had previously shown that mutating the tyrosine residue in the N-Aβcore to a serine [Y10S] or mutating the two histidine residues to two alanines [H13A,H14A] reduces activity, indicating these amino acid residues in the N-Aβcore sequence are essential for activity [13,19]. To confirm the specificity of the N-Aβcore in reversing LTP impairment in 5×FAD hippocampal slices, we tested the reverse-sequence N-Aβcore [QHHVEY] and an inactive triple mutant [SEVAAQ]. Treatment with the reverse-sequence N-Aβcore partially restored LTP in the 5×FAD hippocampal slices (Figs 1D & 1E). Note that the reverse-sequence peptide contains a di-histidine sequence. By contrast, the inactive triple mutant had no significant effect on LTP in the 5×FAD slices (Figs 1D & 1E; 110% ± 13% s.d. compared to 5×FAD). It is important to note that there was no change in basal synaptic transmission or a trend toward increasing LTP in the wild-type slices treated with the reverse N-Aβcore (not shown), as seen for the N-Aβcore (Fig 1). Taken together, these results confirm the contribution of the two essential histidine residues to the positive neuromodulatory activity of the N-Aβcore.

### N-Aβcore rescue of Aβ-induced LTP deficits involves PI3 kinase and mTOR

LTP has shown to involve multiple protein kinase and phosphatase pathways [25,26]. As prior evidence implicates the PI3 kinase and mTOR pathways in the regulation of Aβ neurotoxicity [27,28] and in the regulation of LTP [29], we evaluated the roles of PI3 kinase and mTOR in the action of the N-Aβcore in reversing impaired LTP via treatment of the hippocampal slice preparations with selective inhibitors. As shown in Figure 3, application of PI3 kinase inhibitor LY294002 had no impact on LTP in hippocampal slices from control (background B6.SJL) mice or 5×FAD mice. In contrast, application of LY294002 prior to treatment with the N-Aβcore prevented the rescue by the N-Aβcore of LTP in the slices from 5×FAD mice (Fig 3; 94.3% ± 15% s.d. compared to 5×FAD).

**Fig. 3.**
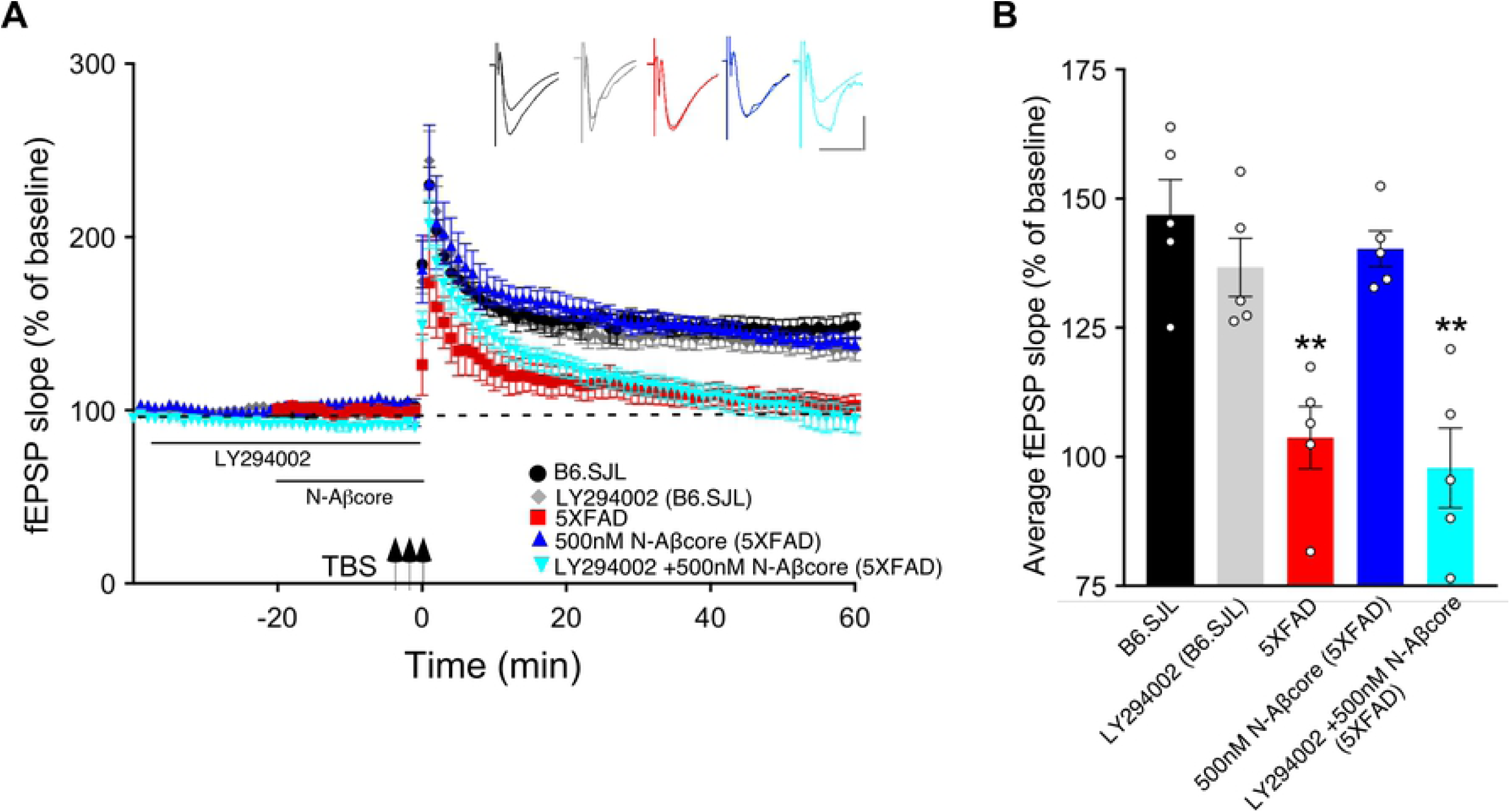
N-Aβcore protection of Aβ-induced synaptic dysfunction involves the PI3 kinase pathway. **A** TBS-induced LTP for B6.SJL slices perfused with control aCSF (black) or 100µM LY294002 (grey) and for 5×FAD slices with aCSF (red), 500nM N-Aβcore (light blue) or 500nM N-Aβcore plus 100µM LY294002 (dark blue). Color-coded inserts showing examples of fEPSPs (baseline vs. LTP). **B** Quantification of average fEPSP slope values for 50-60 min post-TBS (one-way ANOVA: F_(4,20)_= 13.63, p< 0.0001). Bonferroni post hoc tests showed that rescue of the LTP deficit in the 500nM N-Aβcore-treated 5XFAD slices (**p=0.004, 500nM N-Aβcore 5XFAD vs 5XFAD; p<0.05, vs aCSF B6.SJL) was blocked by 100µM LY294002 pretreatment (**p=0.0002, 100µM LY294002 + 500nM N-Aβcore vs. aCSF B6.SJL). Arrows indicate timing of TBS in **A**. All data are expressed as means ± s.e.m. *N*=4 for each condition (1 slice/mouse). Inset calibration: horizontal, 10ms; vertical, 0.4 mV. **p<0.005.

Prior inhibition of the PI3 kinase pathway effector mTOR by rapamycin also prevented the rescue by the N-Aβcore of LTP in the slices from 5×FAD mice (Fig 4; 101% ± 20% s.d. compared to 5×FAD). However, rapamycin did reduce LTP in the control B6.SJL slices, to a level similar to that seen for the LTP in 5×FAD slices. The inhibitor had no significant effect on the reduced level of LTP in the 5×FAD slices.

**Fig. 4.**
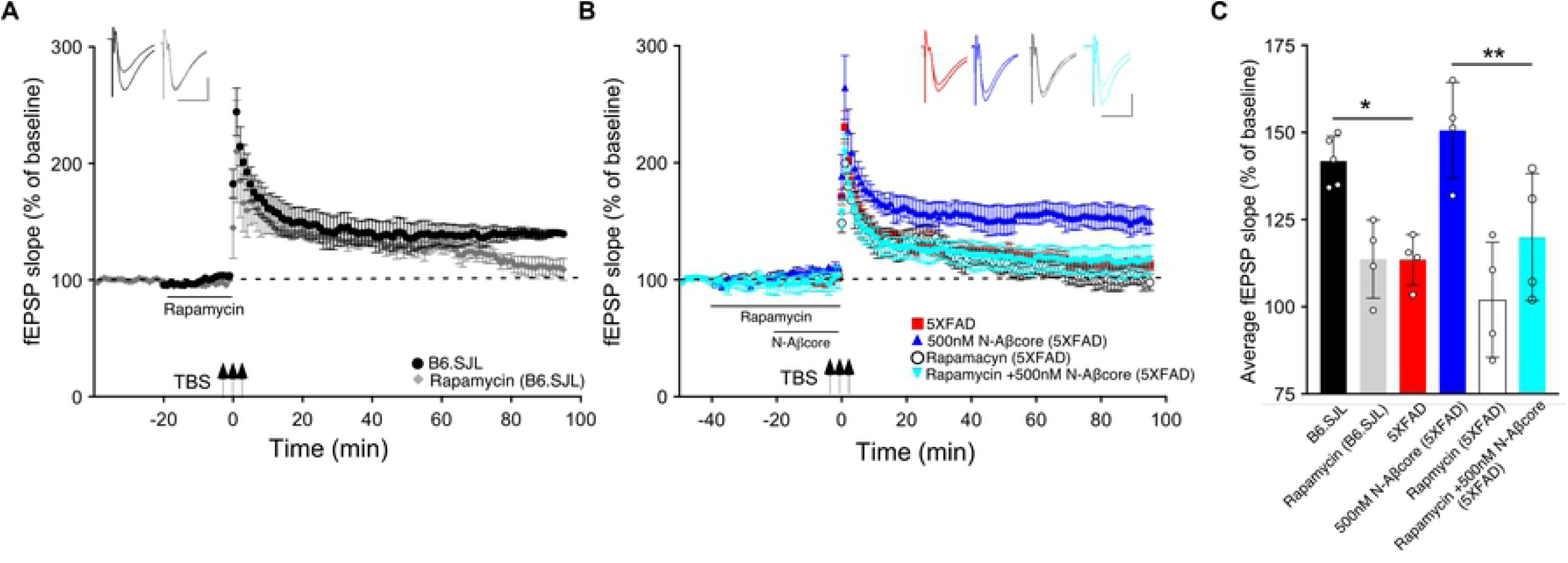
N-Aβcore protection of Aβ-induced synaptic dysfunction involves mTOR downstream in the PI3 kinase pathway. **A** TBS-induced LTP for B6.SJL slices perfused with control aCSF (black) or 500nM rapamycin (grey). **B** TBS-induced LTP for 5×FAD slices with aCSF (red), 500nM rapamycin (white circle), 500nM N-Aβcore (cyan) or 500nM N-Aβcore plus 500nM rapamycin (blue). **C** Quantification of average fEPSP slope values for 50-60 min post-TBS (one-way ANOVA: F_(5,19)_= 8.85, p= 0.0002). B6.SJL slices incubated in 500nM rapamycin (20 min) exhibited reduced LTP when compared to aCSF-treated B6.SJL control slices (Bonferroni post hoc test, *p= 0.038). Rescue of LTP deficit following 20 min 500nM N-Aβcore treatment in 5XFAD slices was abolished with 500nM rapamycin pretreatment (40 min) (**p= 0.0097). Arrows indicate timing of TBS in **A** and **B**. All recording data (**A, B**) are means ± s.e.m. Averaged data for 50-60 min post-tetanus (**c**) are means ± s.d. *N*=4 for each condition (1 slice/mouse). Color-coded insets show examples of fEPSPs (baseline vs. LTP). Inset calibration: horizontal, 10ms; vertical, 0.4 mV *p< 0.05; **p<0.005.

To further probe the mechanism by which the N-Aβcore regulates the PI3 kinase pathway, the impact of the core peptide on Aβ-linked regulation of the PI3 kinase and its downstream effectors Akt and mTOR was investigated using mouse hippocampal neuron cultures and mouse hippocampal slices. Treatment of neuron cultures with full-length Aβ (1-42) was shown to downregulate expression of various PI3 kinase, Akt and mTOR transcripts (Fig 5). Co-treatment with the N-Aβcore alleviated the Aβ-induced downregulation of Akt1 and mTOR transcripts (Fig 5).

**Fig. 5.**
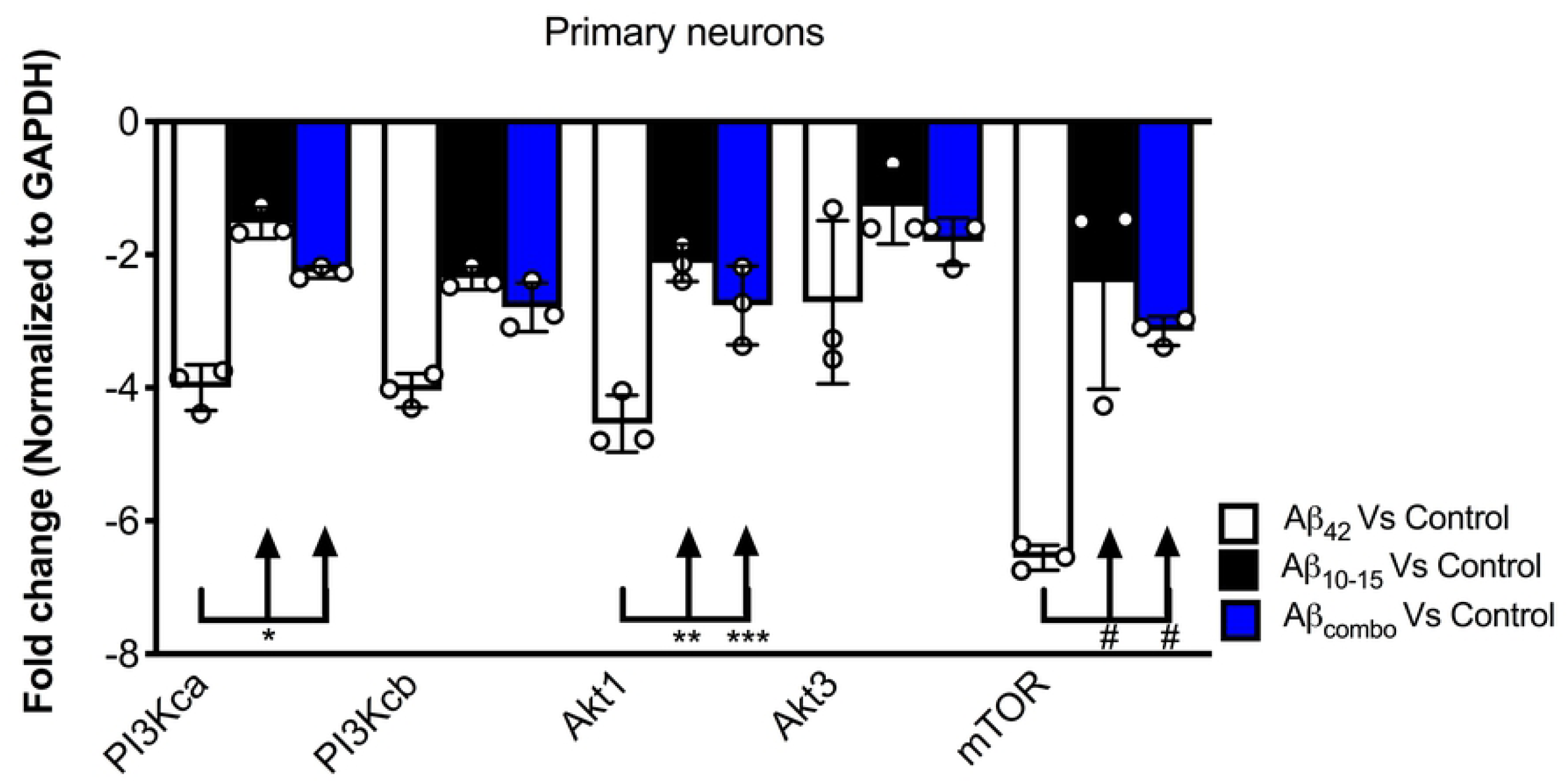
Impact of N-Aβcore on regulation of various PI3 kinase, Akt and mTOR transcripts by Aβ. Primary mouse hippocampal neuron cultures were treated with 1µM full-length Aβ_42_, 1µM Aβ_10-15_ (N-Aβcore) or the combination (combo). Extracted RNA was subjected to qPCR, probing for Class 1 PI3 kinase catalytic subunits (CA and CB), Akt1 and Akt2 and mTOR. Data are expressed as averaged fold-change over untreated controls, after normalizing for GAPDH expression. Bars are means ± s.d. *p= 0.00063; **p= 0.000002; ***p= 0.0001; ^#^p= 0.000001 (Bonferroni post hoc tests for one-way ANOVA comparisons, as shown). For each condition, *N*=3 cultures.

### Elevated levels of Aβ enhances long-term depression and the N-Aβcore reverses Aβ-linked LTD enhancement in hippocampal slices from 5×FAD APP/PS1 mice

LTD is an essential component of synaptic plasticity underlying memory processing in the hippocampus, as synapses cycle between LTP and LTD, a process known as synaptic scaling [30]. To date, few studies have examined the effects of pathological levels of soluble Aβ on LTD induction, and moreover, the results have been mixed. For example, while focusing on NMDA receptor-dependent LTD, administration of synthetic Aβ led to an enhancement of LTD in some cases [eg. 10,31,32], whereas other studies reported no effect [eg. 33]. Here, we aimed to examine the effects of endogenous soluble Aβ on LTD in the 5×FAD hippocampal slices. Using LFS to induce LTP in the same Schaffer collateral – CA1 pathway in hippocampal slice as that used for LTP, the LTD in slices from the 5×FAD mice was more pronounced than that observed for LTD induced in slices from B6.SJL control mice (Figs 6A & 6B; −32.5% ± −5.1% s.d. for 5×FAD compared to −18.1% ± −4.9% s.d for the control B6.SJL slices). Interestingly, treatment of 5×FAD mouse slices with the N-Aβcore prior to and during LFS resulted in a restoration of LTD to the level observed for the B6.SJL slices (Figs 6A & 6B; −21.4% ± 4.5%). The N-Aβcore had no effect on metabotropic glutamate receptor-induced LTD (Fig 7). Taken together, these data suggest that Aβ plays a role in facilitating LTD and the N-Aβcore may protect against Aβ-induced LTD enhancement. The role of the NMDA receptor in Aβ facilitation of LTD warrants further investigation.

**Fig. 6.**
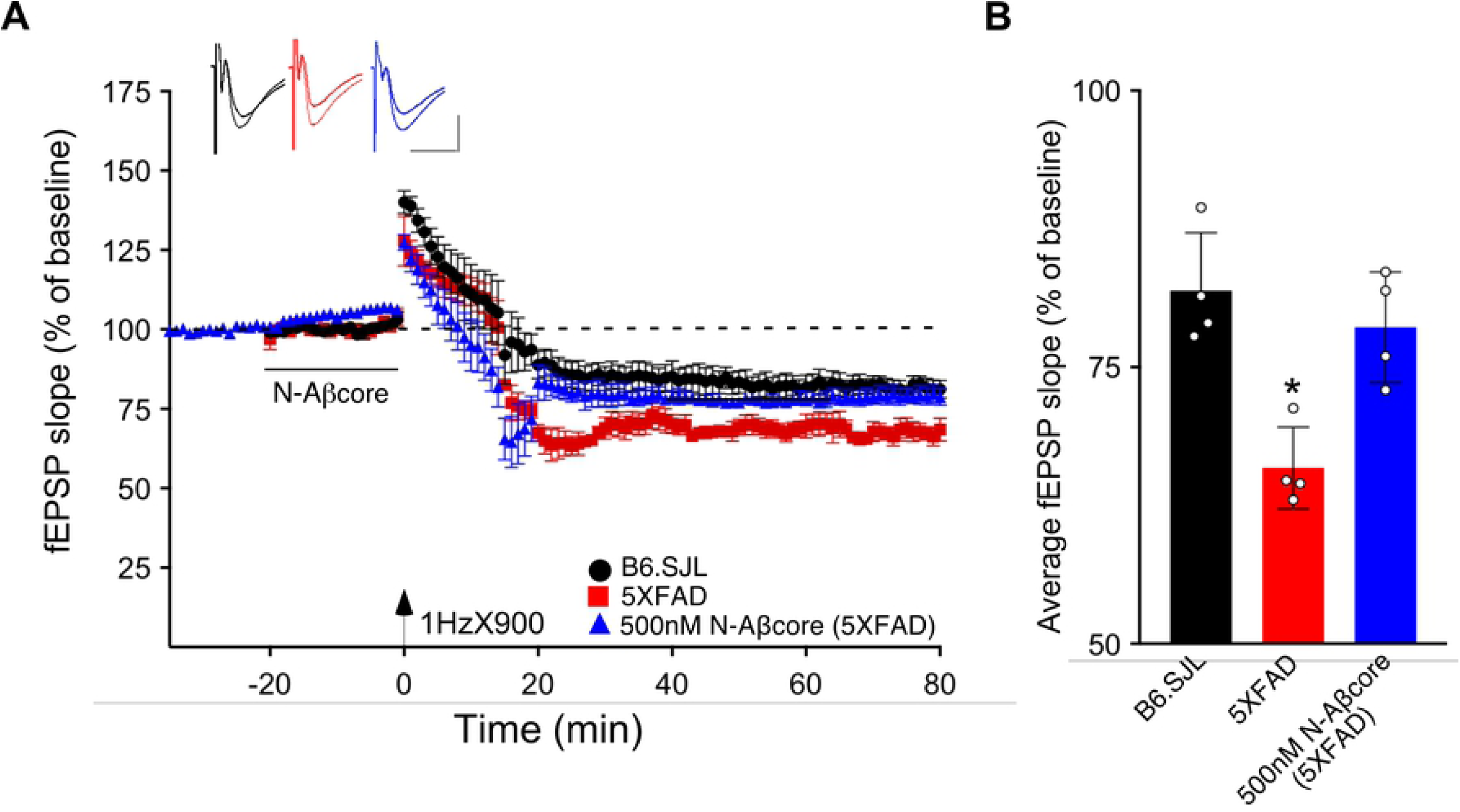
The N-Aβcore reverses endogenous Aβ enhancement of LTD. **A** LFS-induced LTD in hippocampal slices from B6.SJL perfused with control aCSF (black) and 5×FAD slices perfused with aCSF (red) or 500nM N-Aβcore (blue). Color-coded insets show examples of fEPSPs (baseline vs. LTD). **B** Quantification of average fEPSP slope values for 50-60 min post-LFS (one-way ANOVA F_(2,9)_= 13.03, p= 0.0022). Bonferroni post hoc tests: 5×FAD slices displayed significantly more pronounced LTD as compared to control aCSF B6.SJL slices (*p= 0.012). 500nM N-Aβcore in 5XFAD slices returned LTD to the level observed in control aCSF-perfused B6.SJL slices (p>0.99). Data are mean ± s.e.m. *N*=4 for each condition (1 slice/mouse). Inset calibration: horizontal, 10ms; vertical, 0.4 mV. *p< 0.05; **p<0.005.

**Fig. 7.**
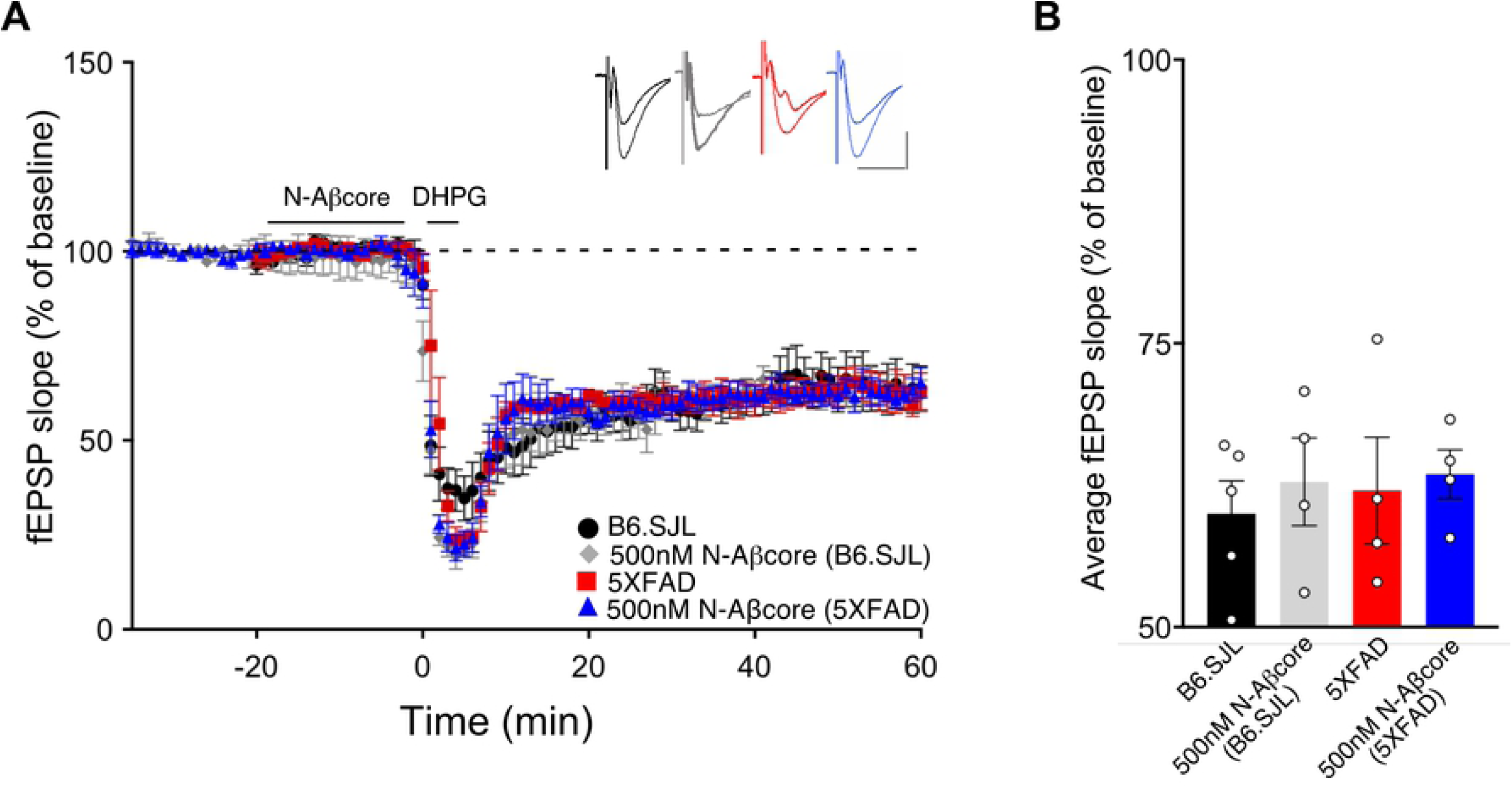
N-AβCore has no impact on NMDA-independent, metabotropic glutamate receptor-induced LTD deficits. NMDA-independent LTD was induced by treatment with bath application of 100µM 3,5-dihydroxyphenylglycine (DHPG), a metabotropic glutamate receptor (mGluR) group I agonist. **A** DHPG-induced LTD in hippocampal slices from B6.SJL mice perfused with control aCSF (black) or 500nM N-Aβcore (grey) and 5×FAD slices perfused with aCSF (red) or 500nM N-Aβcore (blue). Color-coded insets show examples of fEPSPs (baseline vs. LTD). Inset calibration: horizontal, 10ms; vertical, 0.4 mV. **B** Quantification of average fEPSP slope values for 50-60 min post-LFS (p<0.05, two-way ANOVA). Data are means ± s.e.m. *N*=4 for each condition (1 slice/mouse).

## Discussion

Previous studies have established a strong link between the progression of AD and the extent of synaptic dysfunction occurring in the early stages of the disease, prior to the formation of Aβ plaques and tau neurofibrillary tangles [1,5,34,35]. In AD-like models it has been widely demonstrated that elevated levels of soluble oligomeric Aβ drive LTP inhibition [3,7,8,36], coupled to downregulation of synaptic AMPARs. By contrast, the link between pathological levels of Aβ and LTD are less well understood, and as previously noted, investigations of the impact of Aβ on LTD have had conflicting results.

Low “physiological” levels (pM) of soluble Aβ have been shown to enhance synaptic plasticity and facilitate hippocampal-based learning and memory [11-13], suggesting a neuromodulatory role of soluble Aβ at physiological levels. Augmentation of LTP by pM Aβ correlated with enhanced expression of AMPARs. Similar results were obtained using an N-terminal fragment of Aβ (1-15) implicated that sequence within Aβ as accountable for the positive neuromodulatory activity of full-length Aβ [13]. We wondered whether the N-Aβcore (10-15), encompassing the essential sequence within the N-terminal fragment accounting for its positive neuromodulatory activity and its cellular neuroprotective activity against Aβ neurotoxicity, could itself enhance synaptic plasticity and protect against Aβ-induced synaptic dysfunction.

In accordance with previous findings [36], we found that LTP was near absent in the hippocampal slice model from APP/PS1 5×FAD transgenic mice, previously shown to be accounted by elevated Aβ in the brains of the transgenic model mice [20,22]. Treatment here with the N-Aβcore reversed this deficit back to the LTP observed in slices from the background B6.SJL mice. Treatment of 5×FAD slices with the N-Aβcore trended towards LTP enhancement, suggesting that the reversal of the LTP deficits in the 5×FAD slices by the N-Aβcore was not solely due to competitive binding for target receptors and may possibly involve activation of a neuroprotective pathway that enhances synaptic plasticity [see 37]. Here, we identified the PI3 kinase/Akt/mTOR in the reversal of LTP deficits in 5×FAD slices by the N-Aβcore as a primary pathway which has been shown to be a key link to long-term memory [38]. Other downstream pathways engaged by the N-Aβcore are not yet definitely identified, but we suspect that key players involved in synaptic modulation are affected, such as regulation of CREB, PKA, and/or CAMKII or downregulation of calcineurin and/or PP1, subsequently altering AMPA receptor trafficking to the synapses [23,24,39,40], consistent with the observed reversal of downregulation of hippocampal AMPARs by the N-Aβcore in 5×FAD mouse hippocampus. Additionally, the enhancement of the basal synaptic transmission with the treatment of the N-Aβcore suggests a receptor-linked influx of Ca^2+^, which further supports the idea that the N-Aβcore activates an alternative neuroprotective pathway that enhances synaptic plasticity, consistent with results for neuroprotection by the N-Aβcore in Aβ-triggered neurotoxicity [19].

Previously, it has been shown that BDNF enhances basal synaptic transmission [41], therefore, we suspect that the N-Aβcore may be upregulating BDNF expression, possibly through a Ca^2+^-dependent increase in CREB activation and/or expression [42,43]. Another possibility is that the N-Aβcore-induced Ca^2+^ influx could also regulate BDNF release at the synapses, thus, enhancing baseline synaptic transmission and ultimately LTP. It would be interesting to examine the effect of the basal synaptic transmission by the N-Aβcore long-term, and whether the enhancement of LTP observed required changes in baseline transmission prior tetanic stimulation.

In the context of synaptic plasticity, LTD is necessary for neural homeostasis. NMDA receptor-dependent LTD often involves internalization of AMPA receptors via a caspase-dependent pathway [16,44]. To date, however, there is limited understanding in regard to the effects of pathological Aβ on LTD, where some groups show that synthetic Aβ enhances LTD [31,32,45] and others show no effect [33,46], though the differential action of Aβ may have resulted from recording in different subregions of the hippocampus (eg. CA1 vs. dentate gyrus). Here, we found that high concentrations of endogenous soluble Aβ shown to be present in the brains of 5×FAD mice resulted in enhanced LTD in hippocampal slices, and treatment with the N-Aβcore prior to and during the LFS induction of LTD reverses this enhancement. Interestingly, Hu *et. al*. found that applying synthetic soluble Aβ prior to LFS did not affect the early phase of LFS-induced LTD (<2h post LFS), but facilitated the late phase (3-5h post LFS) [45], thus, possibly accounting for different findings. It is important to note that late phase LTP and LTD require new protein synthesis, and mTOR is linked to the regulation of protein synthesis [47]. Indeed, the reversal by the N-Aβcore of LTP deficit in the 5×FAD slices was dependent upon mTOR. As LTD and LTP work in concert to allow for reversible synaptic plasticity and synaptic scaling, the LFS-induced enhancement of LTD in the 5×FAD slices could affect subsequent LTP, and this may be another reason why an LTP deficit was observed in the 5×FAD slices compared to wild-type preparations.

Although NMDA and AMPA receptors are involved in different aspects of LTP and LTD, and their expression is affected by elevated Aβ, metabotropic glutamate receptors (mGluRs) have also been implicated in Aβ-induced synaptic dysfunction [9,45,48,49]. Interestingly, NMDA-independent, mGluR-induced LTD was found to be unaffected by the N-Aβcore. While mGluRs have been linked via cellular prion to Aβ-induced cellular toxicity [49], our findings support a divergence in the Aβ-linked signaling pathways affected by the N-Aβcore in synaptic plasticity. Further work is needed to elucidate the detailed molecular mechanisms involved in N-Aβcore protection or reversal of Aβ-linked synaptic dysfunction, including caspase-dependent intracellular pathways leading to the regulation of AMPA receptors, and eventual synaptic loss in AD.

## Conclusions

The essential core hexapeptide sequence, YEVHHQ, or N-Aβcore, within the neuroprotective N-terminal fragment of Aβ was able to effectively, selectively and potently reverse deficits in synaptic plasticity in hippocampal slices from adult APP/PS1 5×FAD transgenic mice. Attenuated LTP and enhanced LTD observed in the slices from 5×FAD mice were both rescued by the N-Aβcore, returning LTP and LTD back to the levels found for hippocampal slices from background control (B6.SJL) mice. The involvement of the PI3 kinase pathway, specifically, mTOR in the protective action of the N-Aβcore against FAD-linked deficits in synaptic plasticity indicates possible connection between N-Aβcore-linked signaling and the translation machinery at hippocampal synaptic sites (dendritic spines). It would thus be of interest to examine the specific signaling pathways engaged by N-Aβcore at hippocampal sites, especially key downstream elements in translational control such as S6 kinase, a prominent substrate for mTORC1 in neuronal systems.

## Author Contributions

**Conceptualization**: Kelly H. Forest and Robert A. Nichols

**Data curation**: Kelly H. Forest, Ruth Taketa, Komal Arora

**Formal analysis**: Kelly H. Forest, Ruth Taketa, Komal Arora, Cedomir Todorivic

**Funding acquisition**: Robert A. Nichols

**Methodology**: Kelly H. Forest, Ruth Taketa, Komal Arora, Cedomir Todorovic

**Investigation**: Kelly H. Forest, Ruth Taketa, Komal Arora

**Supervision**: Robert A. Nichols

**Validation**: Kelly H. Forest

**Writing – original draft**: Kelly Forest and Robert A. Nichols

**Writing – review & editing**: Kelly Forest, Ruth Taketa, Cedomir Todorovic, Robert A. Nichols

## Acknowledgments

We thank Dr. Naghum Alfulaij for advice. We thank Ms. Ann Hashimoto for mouse breeding and husbandry. We thank Dr. Tessi Sherrin for help with the hippocampal microinjections.

